# Social context and the evolution of delayed reproduction in birds

**DOI:** 10.1101/2023.08.02.551693

**Authors:** Liam U. Taylor, Josef C. Uyeda, Richard O. Prum

**Affiliations:** Department of Ecology & Evolutionary Biology, Yale University, New Haven, CT, United States; Biology Department, Bowdoin College, Brunswick, ME, United States; Department of Biological Sciences, Virginia Polytechnic Institute and State University, Blacksburg, VA, United States

**Keywords:** body size, cooperative breeding, colony nesting, deferred breeding, delayed maturity, fast-slow continuum, immature, juvenile, lek, life history evolution

## Abstract

One puzzling feature of avian life histories is that individuals in many different lineages delay reproduction for several years after they finish growing. Intraspecific field studies suggest that various complex social contexts—such as cooperative breeding groups, nesting colonies, and display leks—result in delayed reproduction because they require forms of sociosexual development that extend beyond physical maturation. Here, we explicitly propose this hypothesis and use a full suite of phylogenetic comparative methods to test it, analyzing the evolution of age at first reproduction (AFR) in females and males across 963 species of birds. Phylogenetic regressions support increased AFR in colonial females and males, cooperatively breeding males, and lekking males. Continuous Ornstein-Uhlenbeck models support distinct evolutionary regimes with increased AFR for all of cooperative, colonial, and lekking lineages. Discrete hidden state Markov models suggest a net increase in delayed reproduction for social lineages, even when accounting for hidden state heterogeneity and the potential reverse influence of AFR on sociality. Our results support the hypothesis that the evolution of social contexts reshapes the dynamics of life history evolution in birds. Comparative analyses of even the most broadly generalizable characters, such as AFR, must reckon with unique, heterogeneous, historical events in the evolution of individual lineages.

## Introduction

In many vertebrates, sexual activity is tightly linked to physical maturity. For example, a House Mouse (*Mus musculus*) reaches full size (∼20g) and begins breeding within a month of birth (Vandenbergh et al. 1972). In contrast, an African Bush Elephant (*Loxodonta africana*) can take decades to reach adult size (>2,000 kg) before breeding (Perry 1953).

Life history theory invokes somatic growth to explain the evolution of delayed reproduction (Williams 1966:87–88; Stearns 1992). Given the ever-present risk of mortality, selection should favor delayed reproduction only if it provides a future reproductive advantage substantial enough to offset the risk of waiting (Wittenberger 1979; Bell 1980; Taylor and Prum 2024). If an increase in body size before maturation contributes to an increase in lifetime fecundity (e.g., a bigger fish can produce more numerous or larger eggs) or a decrease in adult mortality (e.g., a bigger fish is harder to catch and eat), then young organisms may delay reproduction to invest in growth (Roff 1984; Stearns and Koella 1986; Kozłowski 1992).

Extrinsic mortality risks can tune the relative lifetime costs and benefits of juvenile growth, contributing to classic macroevolutionary correlations among large body sizes, long lifespans, and delayed reproduction (e.g., Western and Ssemakula 1982; Harvey and Clutton-Brock 1985; Charnov 1993). A long lifespan, however, is not sufficient to explain delayed reproduction. Selection requires a lifetime fitness benefit from development, such as an increased body size, to outweigh the costs of forgoing reproduction when young (Taylor and Prum 2024).

Birds complicate this logic, because sexual activity is not tightly linked to physical maturity. Nearly all birds grow to adult size within their first year of life, often within weeks or months after hatching (Bennett and Owens 2002). But individuals in many avian species delay reproduction for several years (Lack 1968). If birds do not grow while they delay reproduction, then we cannot understand the evolution of delayed reproduction in terms of physical growth (Charnov 2000). Therefore, although there are strong, macroevolutionary correlations between large body size and delayed reproduction in birds (Western and Ssemakula 1982; de Magalhães et al. 2007), these correlations explain little about the evolution of delayed reproduction in birds.

The alternative hypotheses for avian delayed reproduction highlight behavioral, rather than physical, development. One set of hypotheses is focused on foraging development, suggesting that some birds wait until they develop foraging skills before attempting to rear young (Ashmole 1963; Wunderle 1991). Another explanation—not mutually exclusive with foraging development—is scattered across the literature with regards to avian social behavior (Lack 1968; Orians 1969; Wiley 1974; Gould 1977:345–346; Bradley and Wooller 1990; Zack and Stutchbury 1992; Møller 2006; Hatchwell 2009). Synthesizing these earlier suggestions, we formally propose a hypothesis: some birds have evolved to delay reproduction because they must undergo processes of sociosexual maturation, extending beyond physical maturation, to breed in complex social contexts.

For birds, these complex social contexts include cooperative breeding groups, nesting colonies, and display leks. Cooperative birds have breeding territories that include extra-pair individuals or multi-pair groups (Skutch 1961; Cockburn 2006). Colonial birds defend nest sites, rather than foraging resources, in dense association with conspecific breeders (Perrins and Birkhead 1983). Lekking birds defend display sites, rather than nesting or foraging resources, where they perform elaborate sexual displays in close proximity to others (Bradbury 1981).

Intraspecific field studies suggest these social contexts give rise to delayed reproduction because they create opportunities, or obligations, for sociosexual development. For example, the cooperatively breeding White-winged Chough (Corcoracidae: *Corcorax melanorhamphos*; ∼350g), usually starts breeding around four years old, only after forming a breeding group that assists with incubating and provisioning young (Rowley 1978; Heinsohn and Cockburn 1997). The colonial Snowy Albatross (Diomedeidae: *Diomedea exulans*; ∼7 kg) starts breeding at more than eight years old, only after developing central-place foraging skills, territorial social skills, and pair-bonds, all of which are needed to raise offspring at a pelagic breeding colony (Hector et al. 1986; Weimerskirch and Jouventin 1987; Pickering 1989; Riotte-Lambert and Weimerskirch 2013). Meanwhile, female Long-tailed Manakins (Pipridae: *Chiroxiphia linearis*; ∼20 g) begin breeding in their first or second year, whereas the lekking males spend a minimum of five years—and often more than a decade—navigating the social hierarchies of their cooperative leks and developing coordinated, multi-male sexual displays before they have the chance to copulate (McDonald 1993; Trainer et al. 2002).

Here, we use a broad suite of comparative methods to investigate a fundamental question about avian life histories: by setting the stage for sociosexual development, does the evolution of complex social contexts give rise to the evolution of delayed reproduction? First, we use phylogenetic regressions (Grafen 1989; Garland Jr. et al. 1993) to test overall correlations between social context and age at first reproduction (AFR). Second, we use continuous Ornstein-Uhlenbeck models (Butler and King 2004; Beaulieu et al. 2012) to test whether the mode of AFR evolution differs for lineages living in complex social contexts. Third, we use discrete hidden Markov models (Boyko and Beaulieu 2021) to test whether the evolution of social contexts precedes the evolution of delayed reproduction, or vice versa. Our analyses provide evidence that the historical evolution of social contexts restructures the evolutionary dynamics of delayed reproduction across birds.

## Methods

### Age at first reproduction (AFR)

We coded female and male AFR as an integer value representing the minimum reported breeding age for each species. Our core sources were global and regional encyclopedias, including all full-length articles in Cornell’s Birds of the World (Billerman et al. 2022), The Birds of the Western Palearctic (Cramp et al. 1977), Handbook of Australian, New Zealand, and Antarctic Birds (Marchant and Higgins 1990), and The Birds of Africa (Fry et al. 1982), supplemented with literature on target families or species. We coded separate male and female values whenever possible. If no distinction was made, we used the same value for males and females. References for each species are available in the associated data file.

Most birds in our dataset exhibit some form of annual seasonality, either from temperate winter-summer cycles or tropical wet-dry cycles (Immelmann 1971; Wingfield and Farner 1980; Wikelski et al. 2000). Thus, we summarized AFR as an integer representing annual reproductive season since hatching. For example, an AFR of three indicated that breeding starts in the third annual breeding season, roughly 36 months after hatching.

Criteria for reports of “breeding” differed across taxa, ranging from molecular paternity testing (e.g., male Wire-tailed Manakin *Pipra filicauda*; Ryder et al. 2009), to observations of egg-laying (e.g., female Spanish Eagle *Aquila adalberti*; González et al. 2006), to territory establishment (e.g., male Willow Ptarmigan *Lagopus lagopus*; Hannon and Dobush 1997), to blanket statements about “sexual maturity.” In some cases, information was derived from captive breeding reports (especially for Anseriformes, Galliformes, and Psittaciformes) or plumage maturation timelines for species that breed almost exclusively in definitive plumage (e.g., Regent Bowerbird *Sericulus chrysocephalus*; Frith and Frith 2004). Although varied definitions of “breeding” added interspecific noise to the dataset, most definitions biased AFR downwards (i.e., an individual often needs to hold a territory to build a nest, to lay an egg, to hatch an egg). Our use of minimum, rather than central, AFR values thus offered a more consistent, albeit conservative, representation of breeding age across species.

We did not code intraspecific or intra-annual variation in breeding phenology, which can have important consequences for age-related reproductive outcomes (Becker et al. 2008; López-Calderón et al. 2017; Neate-Clegg and Tingley 2023). One extreme form of intra-annual variation involves birds breeding early in their first year of life, when rapid growth allows for somatic maturity under limited seasonality (e.g., Zebra Finch *Taeniopygia guttata*; Immelmann 1971), across subannual environmental cycles (e.g., *Loxia* crossbills; Hahn 1998), or even within season of hatch (e.g., Anna’s Hummingbird *Calypte anna*; Clark and Russell 2020). There were 37 species in the dataset with a minimum reported AFR of less than one year (Table S1). Further comparative studies would help unpack the relationships among environmental fluctuations, somatic growth, small size, molt, sociosexual development, and life history evolution in these species. In this study, we simply assigned them an AFR of one.

Life history evolution is often more sensitive to changes that occur at younger ages (Hamilton 1966; Williams 1966; Stearns 1992). A shift in younger AFR values (e.g., breeding year two vs. three) may be subject to stronger selection than the same shift in older AFR values (e.g., breeding year eight vs. nine). Therefore, we used log_2_ AFR values for all continuous analyses.

### Social context

We assigned each species in our dataset to one of four social contexts: cooperative, colonial, lekking, or “other.” As introduced previously, cooperative breeding was assigned to species in which breeding territories include extra-pair individuals or groups, encompassing both helpers-at-the-nest (e.g., *Malurus cyaneus*; Dunn et al. 1995) and communal breeding (e.g., *Crotophaga major*; Riehl 2021). Coloniality was assigned to species in which female and male breeding territories exclusively consist of nest sites in close proximity to conspecific breeders (Perrins and Birkhead 1983). Lekking was assigned to species in which one sex has breeding territories that exclusively operate as sexual display sites (Bradbury 1981). This broad definition of lekking (Prum 1994) included species with aggregated “arena” display sites (e.g., *Centrocercus urophasianus*; Wiley 1974) as well as more solitary display sites (e.g., *Ptilonorhynchus violaceus*; Borgia 1985). Species that were neither cooperative, colonial, nor lekking were assigned as social context “other.” As discussed later, this “other” category encompassed a wide range of additional breeding systems and social strategies.

We made minor updates to the Cockburn (2006) parental care dataset for cooperative breeders by including “group” breeders as cooperative (*Cyphorhinus phaeocephalus*, *Nestor meridionalis*, *Nestor notabilis*, *Nymphicus hollandicus*, *Perisoreus canadensis*, *Perisoreus infaustus*, *Pyrrholaemus sagittatus*, and *Zanda funerea*) and by reassigning some species based on more recent life history accounts (*Aphelocoma californica*, *Bucorvus abyssinicus*, *Coragyps atratus*, and *Icterus galbula* were reassigned as not cooperative; *Curruca nisoria*, *Hypsipetes crassirostris*, and *Icterus bullockii* were reassigned as cooperative). We consulted Rolland et al. (1998), Bradbury (1981), and our life history sources for statements about coloniality and lekking. In cases of hierarchically-structured colonies of cooperative breeding groups (e.g., *Manorina melanophrys*; Smith and Robertson 1978), we assigned species under the more proximate context of cooperative. As an additional model covariate, we used mass data (log_10_, not distinguished by sex) from the AVONET dataset (Tobias et al. 2022).

### Phylogenetic trees

We used the distribution of 100 phylogenetic trees from McTavish et al. (2024 [preprint]), pruned to our final dataset (‘random’ node age tree set; https://github.com/McTavishLab/AvesData/commit/1dcd8be23491dce4c8f875699ef6ba203cf54a96). Data and trees were wrangled, pruned, and visualized with the ape, ggtree, and tidyverse packages in R v4.4.1 (Paradis et al. 2004; Yu et al. 2017; Wickham et al. 2019; R Core Team 2024). All analyses were conducted on separate female and male datasets.

### Phylogenetic regressions

First, we directly compared AFR (log_2_) values between social contexts using phylogenetic ANOVAs with Brownian correlation structures from R package phytools (Garland Jr et al. 1993; Revell 2012). If social context influences the evolution of delayed reproduction in birds, we predicted that cooperative, colonial, and lekking lineages should have significantly higher AFR than birds living outside of those social contexts. P-values were simulated from 1,000 iterations for each of 100 trees with post hoc pairwise t-tests. Given three overlapping comparisons per variable (cooperative vs. “other”, colonial vs. “other”, lekking vs. “other”), the Bonferroni-corrected P-value threshold for pairwise tests was α = 0.05/3 = 0.016.

We then tested for overall correlations between AFR (log_2_) and social context (colony, cooperative, lekking, or “other”) using phylogenetic generalized least squares (PGLS) regressions (Grafen 1989). If social context influences the evolution of delayed reproduction in birds, we predicted a significant positive effect on AFR from cooperative breeding, coloniality, and lekking. All models used Brownian correlation structures from R package ape. We fit PGLS models with R package nlme (Pinheiro and Bates 2000) across the full distribution of 100 trees.

We compared three PGLS models for each dataset: Mass-only, Social + Mass, and Social * Mass (Table 1). All models included mass as a covariate. However, mass is not likely a directly confounding covariate with respect to social context; there is not a clear mechanistic hypothesis by which an increase in mass independently influences the evolution of AFR in birds. Instead, we included mass in our models because (1) it is a well-known correlate of AFR in birds and other vertebrates and (2) it serves as a strong proxy for lifespan (Western and Ssemakula 1982; Harvey and Clutton-Brock 1985; Charnov 1993; de Magalhães et al. 2007), which may tune the expected lifetime reproductive benefits of development more generally (Taylor and Prum 2024), and may thus reflect variation in the influence of development operating outside of specific social contexts (e.g., foraging development; Wunderle 1991). In sum, a model effect of mass on AFR (or a model using only mass) offers minimal direct biological insight into AFR evolution for birds, besides correlating with axes of life history variation that are partially orthogonal to social context.

**Table 1.** Phylogenetic regression models for age at first reproduction (AFR) in birds.

| Model<sup>b</sup> | Dataset, $\Delta AIC^a$ | |
| --- | --- | --- |
|  | Female | Male |
| AFR ~ Mass | 31 (0, 62) | 47 (2, 78) |
| AFR ~ Mass + Social | **1 (0, 13)** | **0 (0, 0)** |
| AFR ~ Mass * Social | 11 (1, 26) | 10 (4, 12) |

<sup>a</sup> Median values across 100 trees, with 5–95% quantiles in parentheses. All values rounded.
<sup>b</sup> All AFR values log<sub>2</sub>. All mass values log<sub>10</sub>.

### Continuous evolutionary models

We investigated the influence of social context on AFR evolution in a continuous framework using Ornstein-Uhlenbeck (OU) model comparisons (Butler and King 2004; Beaulieu et al. 2012). If social context influences the evolution of delayed reproduction, then we predicted that more complicated models of evolution—those featuring distinct regimes for cooperative, colonial, and lekking lineages with increased AFR—would outperform simpler models that ignore social context.

We compared seven continuous models (Table 2). These models separated lineages into different regimes for evolution of AFR. Each regime was allowed a distinct central parameter (θ) for AFR. Lineages were assigned a regime based on ancestral state estimates of social context (cooperative, colonial, lekking, or “other”) generated with R package ape (four-state discrete model, symmetrical transition rates). We fit continuous models with R package OUwie (Beaulieu and O’Meara 2022) and compared AIC scores for each of 100 trees.

**Table 2.** Continuous models of age at first reproduction (AFR) evolution in birds, including Brownian motion (BM) and Ornstein-Uhlenbeck (OU) models.

| Model | Regime 1<sup>b</sup> | Regime 2 | Regime 3 | Regime 4 | Dataset, $\Delta AIC^a$ | |
| --- | --- | --- | --- | --- | --- | --- |
|  |  |  |  |  | Female | Male |
| BM | - | - | - | - | 325 (100, 386) | 365 (162, 416) |
| OU1 | All | - | - | - | 47 (18, 63) | 75 (51, 93) |
| OU2-Cooperative | Cooperative | Everything else | - | - | 49 (20, 65) | 76 (52, 94) |
| OU2-Colonial | Colonial | Everything else | - | - | **1 (0, 4)** | 36 (30, 44) |
| OU2-Lekking | Lekking | Everything else | - | - | 47 (16, 63) | 51 (23, 68) |
| OU2-Social | Cooperative, colonial, and lekking | “Other” | - | - | 15 (2, 24) | 33 (24, 41) |
| OU4-Social | Cooperative | Colonial | Lekking | “Other” | **0 (0, 1)** | **0 (0, 0)** |

<sup>a</sup> Median values across 100 trees, with 5–95% quantiles in parentheses. All values rounded.
<sup>b</sup> Models with multiple regimes were assigned distinct central ( $\theta$ ) AFR parameters for each regime

### Discrete evolutionary models

Although AFR resembles a continuous variable, annual seasonality discretizes biological variation to largely integer-scale values. Thus, we also analyzed the evolution of AFR using discrete, hidden state Markov models (Beaulieu et al. 2022). These models investigated not only the correlation between social context and AFR, but also the precedence of these traits in the phylogeny (Pagel 1994) while incorporating background heterogeneity in transition rates (Boyko and Beaulieu 2021). For discrete model comparisons, we predicted that preferred models would show lineages in complex social contexts have elevated transition rates towards increased AFR.

We compared 11 discrete models (Table 3). In these models, AFR was binned into two states (relatively fast reproduction = AFR ≤ 2 vs. relatively slow reproduction = AFR ≥ 3) and sociality was also binned into two states (cooperative, colonial, or lekking vs. “other”). We fit discrete models with R package corHMM (100 random restarts per run; Beaulieu et al. 2022) and compared AIC scores for each of 100 trees.

**Table 3.**
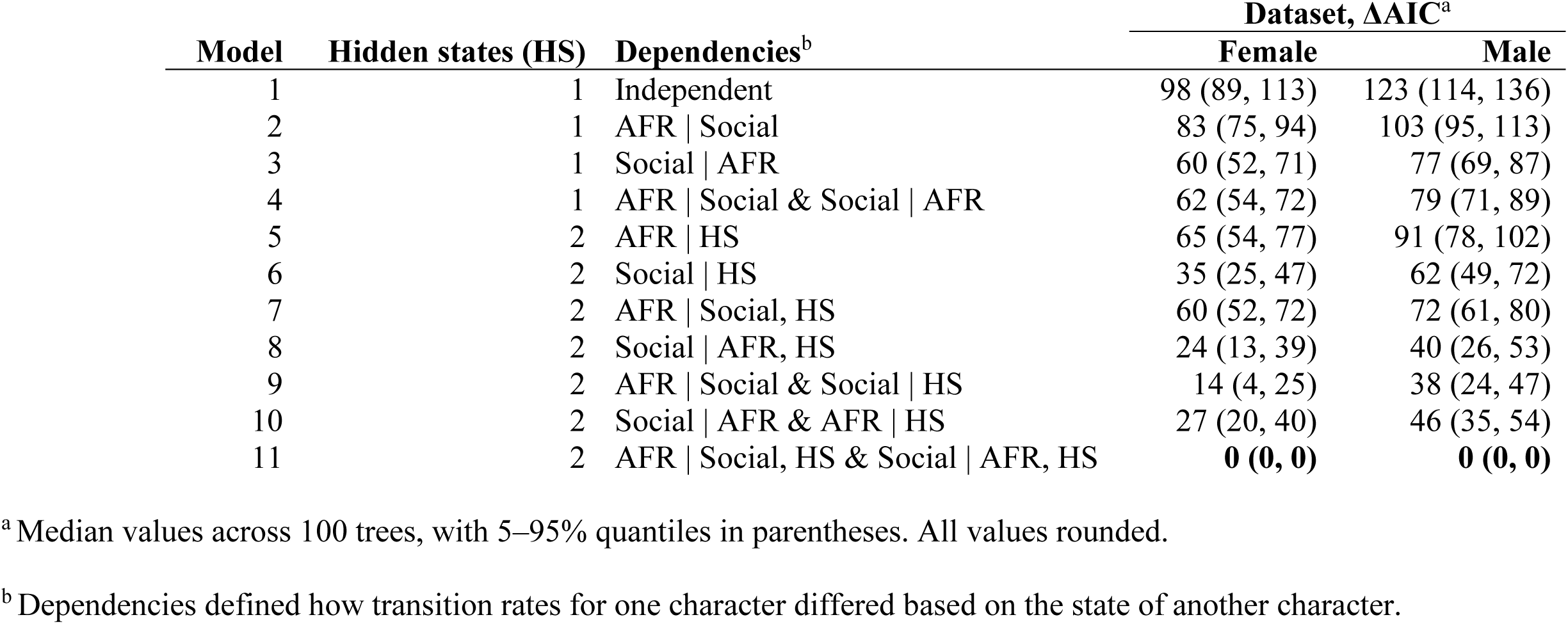
Discrete models of age at first reproduction (AFR ≤ 2 vs. AFR ≥ 3) and social context (cooperative, colonial, or lekking vs. “other”) evolution in birds.

Discrete model parameters are defined in *Supplementary Material: Discrete model descriptions.* Following our hypothesis, a model with elevated transitions towards higher AFR values in the social state should outperform a model in which the evolution of AFR is independent of social state (i.e., model #2 is better than model #1; Table 3). However, this simple comparison neglects two potentially confounding sources of variation. First, a strong association between AFR and sociality may be due to social evolution shifting as a function of AFR, rather than vice versa. We thus included model #3 (in which social transitions depend on AFR state), and model #4 (in which there is a mutual dependence between AFR and social state).

Furthermore, there may be underlying heterogeneity in the evolution of AFR that extends beyond our focal covariate, leading to false preference for any model with multiple AFR parameters (Maddison and FitzJohn 2015; Uyeda et al. 2018; Boyko and Beaulieu 2021). We thus included model #5, which set AFR transitions independent of social state but allowed for heterogeneity via two unobserved hidden states (H1 and H2). Model #6 allowed for hidden state variation in social transitions. More complicated models combined multiple forms of heterogeneity. For example, model #7 let transitions in AFR vary based on both social and hidden states, whereas model #9 let transitions in AFR vary based on social state while sociality, in turn, varied by hidden state. The most complicated model (model #11) featured two hidden states and mutual dependence: in each hidden state, a separate set of parameters described AFR transitions that varied by social state, and, in turn, social transitions that varied by AFR state.

## Results

### Age at first reproduction (AFR)

We compiled minimum reported values of AFR for 963 species (944 species with female values, 950 species with male values, 931 species with either (A) both female and male values or (B) reports that did not distinguish by sex; Fig. 1) across 36/41 orders and 156/251 families in the Clements v2023b taxonomy (Clements et al. 2023). The five orders with no available data were Musophagiformes, Eurypygiformes, Leptosomiformes, Galbuliformes, and Cariamiformes. Of the 95 families with no available data, 69 were in order Passeriformes. The female dataset included 125 cooperative, 199 colonial, and 27 lekking species. The male dataset included 128 cooperative, 200 colonial, and 29 lekking species.

**Figure 1.**
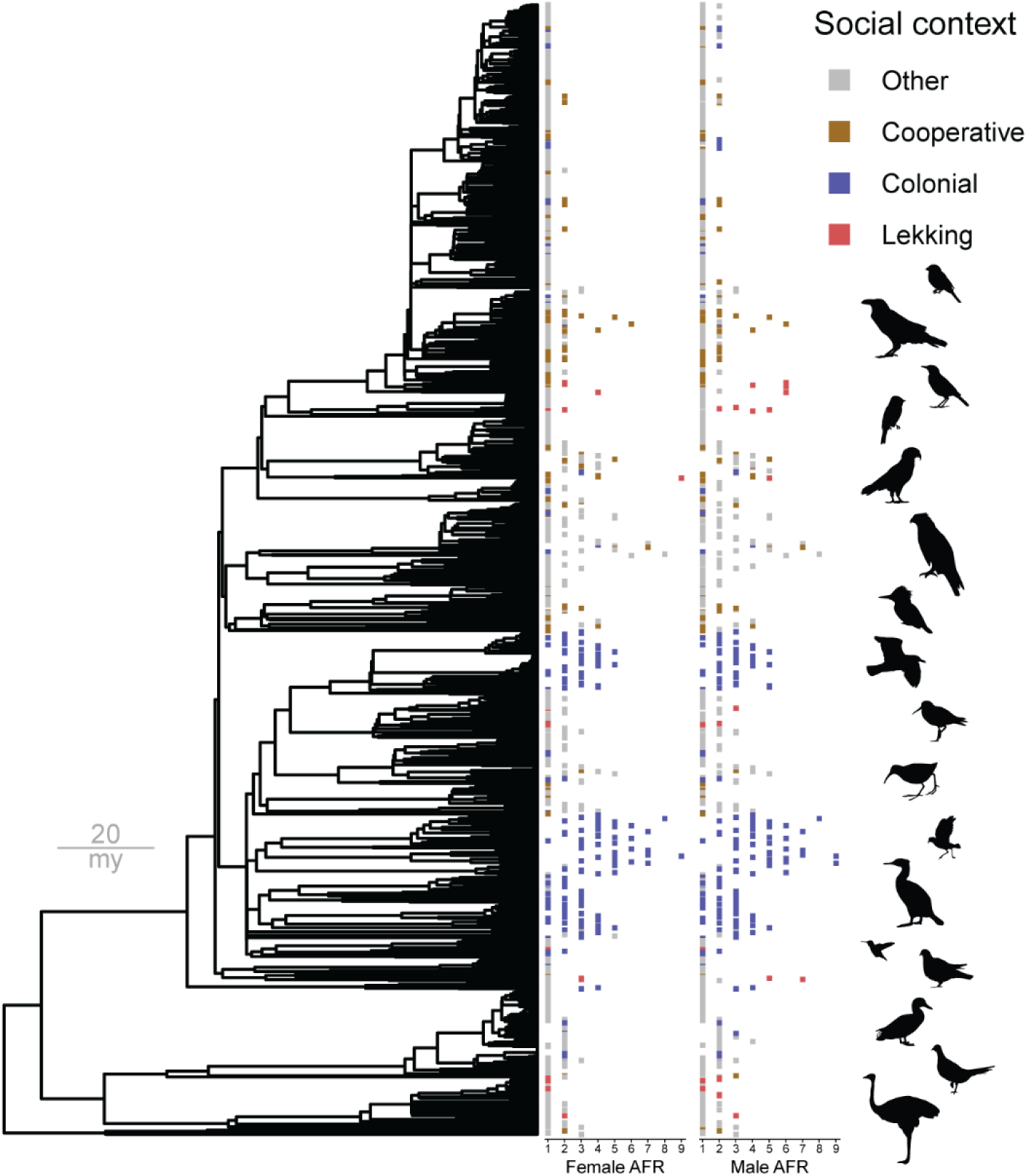
Age at first reproduction (AFR) across the avian phylogeny. Example tree given from distribution of 100 trees from McTavish et al. (2024). Silhouettes from PhyloPic (https://www.phylopic.org/).

Across the dataset, minimum reported AFR values ranged from <1 yr (coded as 1 yr) to 9 yr (Fig. 1). Mean (± SD) AFR for cooperative species was 1.58 yr ± 1.13 for females and 1.61 yr ± 1.14 for males. For colonial species, mean AFR was 2.69 yr ± 1.62 yr for females and 2.74 yr ± 1.64 for males. For lekking species, mean AFR was 1.70 yr ± 1.66 for females, and 2.93 yr ± 2.00 for males. The remaining species, classified as social context “other,” had mean AFR 1.36 ± 0.84 for females and 1.4 ± 0.86 for males.

### Regression analyses

Phylogenetic ANOVAs for AFR across social contexts were overall significant for female, male, and male vs. female difference values (median *P* [5–95% quantiles], female: 0.003 [0.001, 0.006]; male: 0.001 [0.001, 0.003]; male vs. female difference: 0.009 [0.003, 0.017]). Post hoc pairwise t-tests showed colonial birds had significantly increased AFR when compared to “other” species for both females and males (colonial vs. “other”, female: *T* = –15.6, *P* = 0.002 [0.001, 0.004]; male: *T* = –15.5, *P* = 0.001 [0.001, 0.004]). AFR was not significantly different between cooperative and “other” species (cooperative vs. “other”, female: T = –2.2, P = 0.43 [0.34, 0.47]; male: T = –1.9, P = 0.48 [0.43, 0.52]). Lekking birds had increased AFR in males, but not females, although the limited number of lekking birds meant this pairwise comparison was not significant under the conservative Bonferroni-corrected threshold (lekking vs. “other”, female: *T* = –1.2, *P* = 0.70 [0.67, 0.72], male: *T* = –6.6, *P* = 0.02 [0.01, 0.03]). However, lekking birds did show significantly larger sexual differences in AFR (male AFR > female AFR) when compared to “other” species (difference in male vs. female AFR for lekking vs. “other”: *T* = – 13.7, *P* = 0.001 [0.001, 0.001]). In contrast, neither cooperative nor colonial birds showed sexual differences in AFR when compared to “other” species (male vs. female difference in AFR for cooperative vs. “other”: *T* = –0.2, *P* = 0.96 [0.94, 0.97]; colonial vs. “other”: *T* = –0.7, *P* = 0.90 [0.89, 0.92]).

Turning to PGLS models, there was overall preference for the Mass + Social model over the Mass-only or Social * Mass interaction models for both female and male datasets (Table 1). However, variation in relative AIC values across different tree samples for the female dataset (Table 1) suggests that standing phylogenetic uncertainty has substantial consequences on the correlative signals among mass, social context, and female AFR.

All PGLS model estimates are provided in Table S2. Results from the overall preferred Social + Mass model are summarized in Table 4. Across trees, there were universally significant positive effects of mass on both female and male AFR. The significance of cooperative breeding varied, with median P-values of 0.02 for males but 0.055 for females, in both cases ranging to insignificant for some trees. Coloniality had significant positive effects for both females and males, whereas lekking had a significant positive effect for males only (Table 4).

**Table 4.**
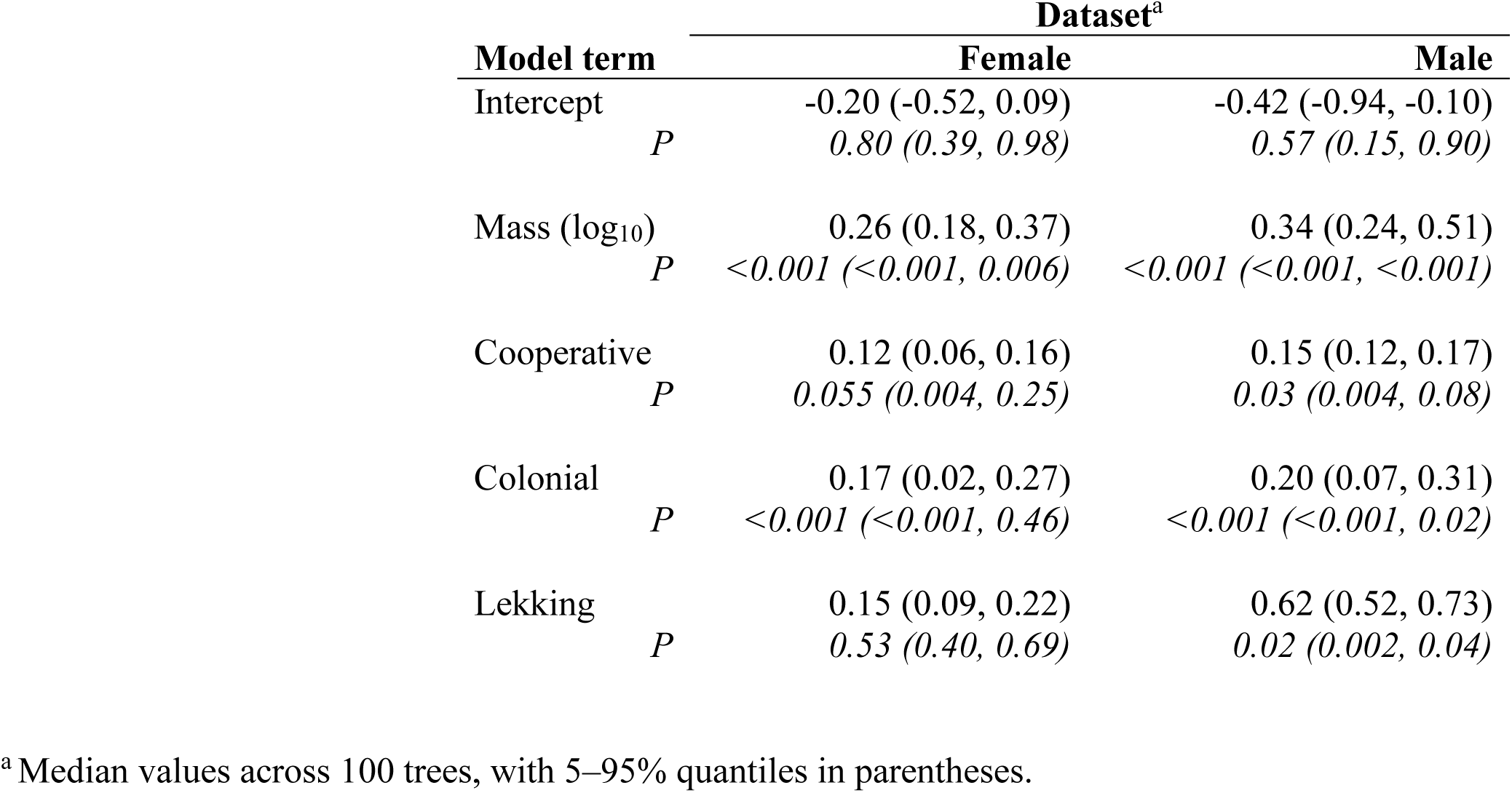
Phylogenetic regression estimates for age at first reproduction (AFR, log_2_) in birds (Mass + Social model)

For a conservative interpretation of model effects, consider a lineage with AFR = 1 yr (2^0^). The log_10_-transformed mass effect of 0.26 (for females) and 0.34 (for males) indicated that a ten-fold increase in mass is associated with an increased AFR of 1.20 yr (2^0.26^) for females and 1.27 yr (2^0.34^) for males (Table 4). In parallel, we can interpret the effects of cooperative breeding with increased AFR of 1.09 yr (2^0.12^) for females and 1.11 yr (2^0.15^) for males, only the latter of which was significant across trees. Coloniality was associated with an increased AFR of 1.12 (2^0.17^) for females and 1.15 (2^0.20^) for males. The larger lekking effect for males would correspond to an increased AFR of 1.53 (2^0.62^).

Another way to interpret these correlations is to directly compare social and mass effects with one another. A ten-fold increase in mass (i.e., one unit increase in log_10_ mass = 10^1^) had an estimated effect of ∼0.30 (Table 4). Cooperative and colonial effect sizes were ∼0.15, or roughly 50% the magnitude of the mass effect (i.e., an effect equivalent to half a unit increase in log_10_-transformed mass). In other words, the effects of cooperative breeding and coloniality on AFR were roughly equivalent to a tripling of mass (10^0.5^ = 3.16). The male lekking effect (0.62) was roughly twice the size of the male mass effect (0.34), meaning the effect of lekking on male AFR was roughly equivalent to a hundred-fold increase in mass.

### Continuous evolutionary models

When comparing continuous models of AFR evolution, all OU models outperformed a BM model for both female and male datasets (Table 2). For the male dataset, the OU4-Social model was clearly preferred, with four distinct regimes for each of cooperative, colonial, lekking, and “other” lineages (Table 2). For the female dataset, the OU4-Social model was marginally preferred to the OU2-Colonial model across trees, the latter of which only distinguished colonial species in a separate regime (Table 2). These two models, OU4-Social and OU2-Colonial, outperformed all others for the female dataset (Table 2).

All continuous model estimates are provided in Table S3. Results from the clearly preferred (for males) or marginally preferred (for females) OU4-Social model are summarized in Table 5. The log_2_-transformed median θ values for the “other” regime corresponded to an AFR of 1.3 yr for females and males. Estimated θ values for cooperative regimes were only slightly elevated, at 1.5 yr for female and males, whereas colonial θ values corresponded to 2.4 yr.

**Table 5.**
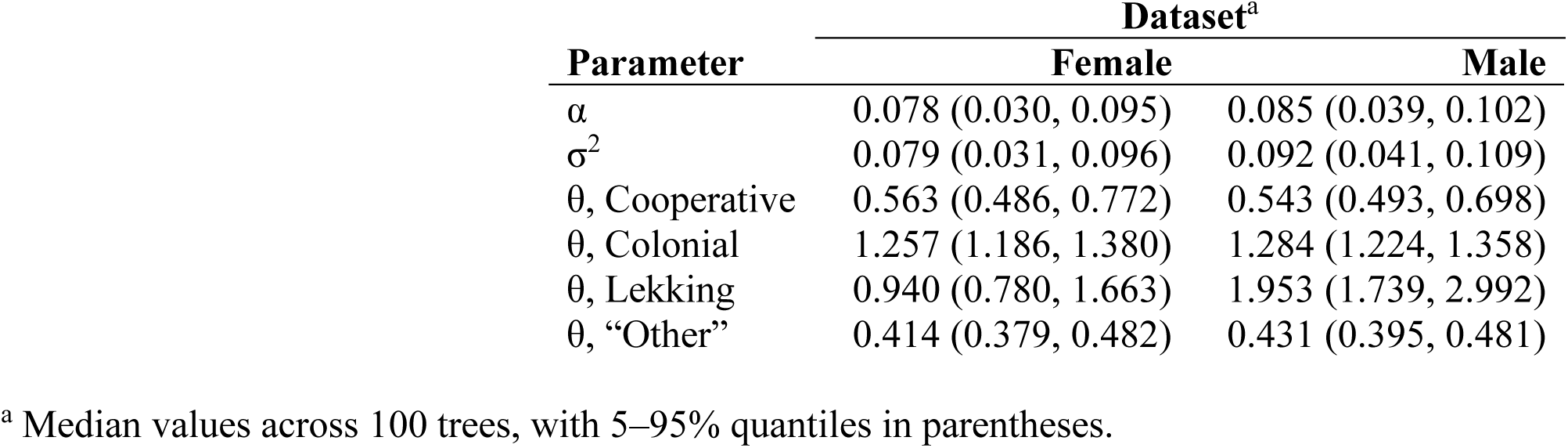
Continuous evolutionary model estimates for age at first reproduction (AFR, log_2_) in birds (OU4-Separate model)

Estimated θ for the lekking regime was more than twice as high in males than in females, corresponding to 1.9 yr for females and 3.9 yr for males. Estimated α parameters—which were shared across all regimes—indicated a phylogenetic half-life of 8.9 Myr and 8.2 Myr for females and males, respectively, out of a total tree depth ranging 72–133 Myr across the full distribution of 100 trees.

Estimated θ values were increased in colonial, lekking, and combined social regimes for every model in which those parameters were estimated (Table S3). In the OU2-Cooperative model, θ for the cooperative regime was lower relative to the regime combining colonial, lekking, and “other” (Table S3). The OU2-Colonial model for females, which rivalled the OU4-Social model in terms of AIC (Table 2), gave log_2_-transformed θ values corresponding to AFR of 2.4 yr for the colonial regime and 1.4 yr for everything else (Table S3).

### Discrete evolutionary models

All discrete model estimates are provided in *Supplementary material: Discrete model descriptions*. The best available model for both female and male datasets was the most complex (model #11), involving hidden state heterogeneity in both sociality and AFR, as well as mutual dependence in those two variables (Table 3). Generally, parameter estimates from model #11 supported the hypothesis that evolution from fast breeding (i.e., AFR ≤ 2) to slow breeding (i.e., AFR ≥ 3) occurred more frequently in lineages living in complex social contexts, even when allowing for mutual dependency and hidden state heterogeneity.

Model #11 included two hidden states (H1 and H2) that influenced transitions in both AFR and social context. The values of H1 and H2 are unrelated when comparing male vs. female results. In female H1, transitions from fast to slow breeding occurred at twice the rate in social lineages (0.0504 for social vs. 0.0243 for “other”) and the opposite transition, from slow to fast breeding, also occurred an order of magnitude less in social lineages (0.0188 for social vs. 0.3354 for “other”). Surprisingly, in female H2, transitions from fast to slow breeding were less frequent in social lineages (<0.0001 for social vs. 0.0008 for “other”). But this difference was offset by an even greater difference in the opposite direction, from slow to fast breeding (0.0057 for social vs. 0.0361 for “other”). Similarly, male H1 had more frequent transitions to slow breeding in social lineages (0.0019 for social vs. <0.0001 for “other”) and less frequent transitions to fast breeding in social lineages (0.0247 for social vs. 0.0305 for “other”). Male H2 showed a remarkable decrease in transitions from fast to slow breeding in social lineages (0.0707 for social vs. 0.4137 for “other”) but this was again offset by an even greater difference in the opposite direction, from slow to fast breeding (0.0268 for social vs. 11.8766 for “other”).

Interpretation of model #11 is complicated by the large number of parameters and their wide estimation intervals. Our overall suite of discrete models, however, supports claims that (1) AFR evolution partially depends on social state while (2) social state is evolving with considerable underlying heterogeneity. When comparing discrete models with no hidden states, there was marginal preference for a model in which social state was dependent on AFR (model #3) versus a model with mutual dependence between sociality and AFR (model #4) for both female and male datasets (Table 3). But both models were far outperformed by the overall second-best model (#9), in which AFR depends on social state, while social state depends on hidden state (Table 3). Thus, performance of model #3 may largely be attributed to underlying heterogeneity in social evolution.

In model #9, transitions in social state were binned into one hidden state with rapid transitions (in both directions) and a second hidden state that was nearly static, with wide uncertainty in those parameters across trees (*Supplementary material: Discrete model descriptions*). Meanwhile, transitions from fast to slow breeding were higher in social lineages compared to “other” lineages (32.3 times higher for females, 18.3 times higher for males).

Transitions from slow to fast breeding were also lower in social lineages compared to “other” lineages (2.8 times lower for females, 2.3 times lower for males; *Supplementary material: Discrete model descriptions*). We caution that strong support for model #11 suggests considerably more complexity—potentially arising from context-dependence, confounding factors, or hidden structure in the binary traits examined.

## Discussion

Overall, our battery of comparative analyses supports the hypothesis that alternative social contexts—cooperative breeding, coloniality, and lekking—give rise to the evolution of delayed reproduction across the avian phylogeny (Fig. 1). Phylogenetic ANOVAs revealed a significant increase in age at first reproduction (AFR) for colonial birds in both male and female datasets. PGLS models, incorporating mass as a proxy for additional covariates, revealed significant increases in AFR for colonial males and females, along with males of cooperative and lekking species (Tables 1, 4). Continuous evolutionary models identified distinct regimes for cooperative, colonial, and lekking lineages, each exhibiting increased AFR in both sexes (Tables 2, 5). Discrete evolutionary models suggested that AFR evolution is influenced by social state, with lineages in complex social contexts experiencing a net shift toward delayed reproduction (*Supplementary material: Discrete model descriptions*). These results provide quantitative, comparative support for earlier conjectures that sociality influences life history evolution across birds (Wiley 1974; Gould 1977; Bradley and Wooller 1990; Zack and Stutchbury 1992; Møller 2006).

Intraspecific field studies reveal the different ways in which social contexts mechanistically influence delayed reproduction through the demands of sociosexual development. For example, colonial seabirds must first gain social experience with nesting territories (Ainley et al. 1983; Taylor 2024) and establish a pair-bond (Pickering 1989) before breeding among conspecifics in dense island colonies. Since most colonial birds are socially monogamous (Lack 1968; Cockburn 2006), these developmental demands apply to young males and young females alike. In contrast, lekking species show a pronounced asymmetry in the demands of sociosexual maturation. Young males, but not young females, must establish display sites, courtship displays, and relationships with other males before they can reproduce (e.g., McDonald 1989; Collis and Borgia 1993; Prum 2017).

The sexual asymmetries of lekking are clear in our results, consistent with previous comparative studies (Wiley 1974; Ancona et al. 2020). The estimated PGLS effect of lekking on log_2_ AFR was much higher for males than females (0.62 in males vs. 0.15 for females; Table 4). The central OU parameters for log_2_ AFR were much higher for the lekking regime in males than females (1.95 in males vs. 0.94 in females; Table 5). Phylogenetic ANOVAs supported a significant elevation in male AFR—relative to female AFR—for lekking lineages, but not colonial or cooperative lineages.

Social behavior shapes avian life histories. This principle extends beyond birds. Many studies demonstrate how sociosexual development influences the life histories of mammals and fish (e.g., Robertson 1972; Gould 1977:345–351; Rodd et al. 1997; Holekamp and Strauss 2020; Kralick et al. 2023). Indeed, our opening caricature of AFR-mass relationships in mice and elephants is also a reference to cases where social environment influences sexual maturity. The presence of males speeds maturation in young female House Mice (Vandenbergh et al. 1972) and suppresses maturation in young male African Elephants (Slotow et al. 2000).

However, our results also highlight significant heterogeneity in avian life history evolution. PGLS models included a significant relationship between mass and AFR (Tables 1, 5). Since mass itself cannot mechanistically explain variation in AFR across birds (Charnov 2000), the significant effect of mass suggests that hidden covariates underlie additional variation in AFR. Indeed, discrete evolutionary models that allowed for hidden state heterogeneity—whether for sociality, or AFR, or both—outperformed related models that did not include such heterogeneity (e.g., model #7 vs. #2, #11 vs. #4; Table 3).

One source of heterogeneity is the fact that variation in avian social complexity is not fully captured by our simple categories of cooperative breeding, coloniality, lekking, and “other.” In particular, our “other” category neglected a huge diversity of alternative avian societies. Examples include complex social networks in parrots (Penndorf et al. 2023) and dominance hierarchies in scavenging raptors (Dwyer and Cockwell 2011). The Black Vulture (*Coragyps atratus*)—one scavenger we coded as social context “other”—has elaborate, communal roosting behavior and does not start breeding until eight years old (Parker et al. 1995).

Another limit to our analysis involved intraspecific variation. We relied on minimum reported AFR to help maintain consistency across species, but this approach overlooks variation within species, which itself may be socially mediated. Intraspecific variation is especially pronounced in cooperative breeders, where territory availability and environmental fluctuations (e.g., rainfall) can influence decisions about helping, dispersing, or breeding (Koenig et al. 1992; Rubenstein 2011; Welklin et al. 2023). For instance, the cooperative Mexican Jay (*Aphelocoma wollweberi*) has a minimum AFR of two years—the value used in our analysis—but some individuals delay reproduction until 14 years old (McCormack and Brown 2020). Our simple trait categorizations thus neglect well-known variation in the life histories of birds.

A further source of variation known to influence avian delayed reproduction is foraging development (Ashmole 1963; Wunderle 1991). There may be no straightforward way to separate the impacts of foraging and social development; foraging is deeply linked to social context in multiple different ways across the avian phylogeny. For example, colonial seabirds must develop central-place foraging skills because they breed at remote island colonies (Coulson 2001; Collet et al. 2020). Snow (1971) proposed that frugivory contributes to the evolution of polygynous leks by making it easier for a single parent to raise offspring. Wiley (1978) proposed that lekking evolves in highly precocial lineages, such as grouse, because one parent can support hatchlings that feed themselves. Thus, for lekking birds, unlike colonial seabirds, it may be the *ease* of foraging that leads to sex-specific delays in reproduction.

Ultimately, our study suggests that the ecological and social diversity of birds confounds the search for simple, lawlike generalizations about their life histories (Taylor and Prum 2024). Life history variables might appear universal; every species might have some age at first reproduction, some clutch size, some lifespan, and so on. A mainstream tradition searches for simple, ahistorical correlations among these life history variables, treating phylogenetic history itself as statistical noise along the way (e.g., Western and Ssemakula 1982; Harvey and Clutton-Brock 1985; Adler et al. 2014; Salguero-Gómez et al. 2016; Healy et al. 2019; see Stearns 1983 for the roots of an alternative approach).

Understanding avian evolution requires more nuance. On one hand, our results support a general pattern: the evolution of social complexity leads to the evolution of delayed reproduction. On the other hand, social complexity itself emerges from individual evolutionary events—the origins of lekking here, the origins of coloniality there—the histories of which fall outside the scope of our general pattern. Thus, even for an apparently universal character such as age at first reproduction, unique events in phylogenetic history remain key to understanding evolution (Uyeda et al. 2018).

## Acknowledgements

We thank Martha Muñoz, Stephen Stearns, Casey Dunn, and Henry Camarillo for helpful comments on the manuscript. This work was supported by an NSF GRFP (#DGE1752134) for L.U.T. and the W.R. Coe fund from Yale University. We are grateful to the Yale Center for Research Computing (McCleary HPC) and Bowdoin College Information Technology for guidance and access to research computing infrastructure.

## References

1. Adler P.B., Salguero-Gómez R., Compagnoni A., Hsu J.S., Ray-Mukherjee J., Mbeau-Ache C., Franco M. 2014. Functional traits explain variation in plant life history strategies. Proceedings of the National Academy of Sciences. 111:740–745.

2. Ainley D.G., LeResche R.E., Sladen W.J. 1983. Breeding biology of the Adélie Penguin. Berkeley: University of California Press.

3. Ancona S., Liker A., Carmona-Isunza M.C., Székely T. 2020. Sex differences in age-to-maturation relate to sexual selection and adult sex ratios in birds. Evolution Letters. 4:44– 53.

4. Ashmole N.P. 1963. The regulation of numbers of tropical oceanic birds. Ibis. 103:458–473.

5. Beaulieu J.M, O’Meara B., Oliver J., Boyko J. 2022. corHMM: Hidden Markov Models of Character Evolution. R package version 2.8. https://CRAN.R-project.org/package=corHMM.

6. Beaulieu J.M., Jhwueng D.-C., Boettiger C., O’Meara B.C. 2012. Modeling stabilizing selection: Expanding the Ornstein–Uhlenbeck model of adaptive evolution. Evolution. 66:2369– 2383.

7. Beaulieu J.M., O’Meara B. 2022. OUwie: Analysis of evolutionary rates in an OU framework. R package version 2.10. https://CRAN.R-project.org/package=OUwie.

8. Becker P.H., Dittmann T., Ludwigs J.-D., Limmer B., Ludwig S.C., Bauch C., Braasch A., Wendeln H. 2008. Timing of initial arrival at the breeding site predicts age at first reproduction in a long-lived migratory bird. Proceedings of the National Academy of Sciences. 105:12349–12352.

9. Bell G. 1980. The costs of reproduction and their consequences. The American Naturalist. 116:45–76.

10. Bennett P.M., Owens I.P. 2002. Evolutionary ecology of birds: Life histories, mating systems and extinction. New York: Oxford University Press.

11. Billerman S.M., Keeney B.K., Rodewald P.G., Schulenberg T.S, editors. 2022. Birds of the World. Ithaca: Cornell Lab of Ornithology.

12. Borgia G. 1985. Bower quality, number of decorations and mating success of male Satin Bowerbirds (*Ptilonorhynchus violaceus*): An experimental analysis. Animal Behaviour. 33:266–271.

13. Boyko J.D., Beaulieu J.M. 2021. Generalized hidden Markov models for phylogenetic comparative datasets. Methods in Ecology and Evolution. 12:468–478.

14. Bradbury J.W. 1981. The evolution of leks. In: Alexander R.D., Tinkle D.W., editors. Natural selection and social behavior. New York: Chiron Press. p. 138–169.

15. Bradley J.S., Wooller R.D. 1990. Philopatry and age of first-breeding in long-lived birds. Acta XX Congressus Internationalis Ornithologici. 3:1657–1665.

16. Butler M.A., King A.A. 2004. Phylogenetic comparative analysis: A modeling approach for adaptive evolution. The American Naturalist. 164:683–695.

17. Charnov E.L. 1993. Life history invariants: Some explorations of symmetry in evolutionary ecology. Oxford: Oxford University Press.

18. Charnov E.L. 2000. Evolution of life-history variation among species of altricial birds. Evolutionary Ecology Research. 2:375–383.

19. Clark C.J., Russell S.M. 2020. Anna’s Hummingbird (*Calypte anna*). In: Poole A.F., editor. Birds of the world. Ithaca: Cornell Lab of Ornithology.

20. Clements J.F., Rasmussen P.C., Schulenberg T.S., Iliff M.J., Fredericks T.A., Gerbracht J.A., Lepage D., Spencer A., Billerman S.M., Sullivan B.L., Wood C.L. 2023. The eBird/Clements checklist of birds of the world: v2023b. Ithaca: Cornell Lab of Ornithology. https://www.birds.cornell.edu/clementschecklist/download/.

21. Cockburn A. 2006. Prevalence of different modes of parental care in birds. Proceedings of the Royal Society B: Biological Sciences. 273:1375–1383.

22. Collet J., Prudor A., Corbeau A., Mendez L., Weimerskirch H. 2020. First explorations: Ontogeny of central place foraging directions in two tropical seabirds. Behavioral Ecology. 31:815–825.

23. Collis K., Borgia G. 1993. The costs of male display and delayed plumage maturation in the Satin Bowerbird (*Ptilonorhynchus violaceus*). Ethology. 94:59–71.

24. Coulson J.C. 2001. Colonial breeding in seabirds. In: Schreiber E.A., Burger J., editors. Biology of marine birds. Boca Raton: CRC Press. p. 100–127.

25. Cramp S., Brooks D., Perrins C. 1977. Handbook of the birds of Europe, the Middle East, and North Africa: The birds of the western palearctic. Oxford: Oxford University Press.

26. Dunn P.O., Cockburn A., Mulder R.A. 1995. Fairy-wren helpers often care for young to which they are unrelated. Proceedings of the Royal Society B: Biological Sciences. 259:339– 343.

27. Dwyer J.F., Cockwell S.G. 2011. Social hierarchy of scavenging raptors on the Falkland Islands, Malvinas. Journal of Raptor Research. 45:229–235.

28. Frith C.B., Frith D.W. 2004. Bowerbirds. Oxford: Oxford University Press.

29. Fry C.H., Urban E.K., Keith S., Safford R., Hawkins F. 1982. The birds of Africa. Cambridge: Academic Press.

30. Garland Jr T., Dickerman A.W., Janis C.M., Jones J.A. 1993. Phylogenetic analysis of covariance by computer simulation. Systematic Biology. 42:265–292.

31. González L.M., Oria J., Margalida A., Sánchez R., Prada L., Caldera J., Aranda A., Molina J.I. 2006. Effective natal dispersal and age of maturity in the threatened Spanish Imperial Eagle *Aquila adalberti*: Conservation implications. Bird Study. 53:285–293.

32. Gould S.J. 1977. Ontogeny and phylogeny. Cambridge: Harvard University Press.

33. Grafen A. 1989. The phylogenetic regression. Philosophical Transactions of the Royal Society B: Biological Sciences. 326:119–157.

34. Hahn T.P. 1998. Reproductive seasonality in an opportunistic breeder, the Red Crossbill, *Loxia curvirostra*. Ecology. 79:2365–2375.

35. Hamilton W.D. 1966. The moulding of senescence by natural selection. Journal of Theoretical Biology. 12:12–45.

36. Hannon S.J., Dobush G. 1997. Pairing status of male Willow Ptarmigan: Is polygyny costly to males? Animal Behaviour. 53:369–380.

37. Harvey P.H., Clutton-Brock T.H. 1985. Life history variation in primates. Evolution. 39:559– 581.

38. Hatchwell B.J. 2009. The evolution of cooperative breeding in birds: Kinship, dispersal and life history. Philosophical Transactions of the Royal Society B: Biological Sciences. 364:3217–3227.

39. Healy K., Ezard T.H.G., Jones O.R., Salguero-Gómez R., Buckley Y.M. 2019. Animal life history is shaped by the pace of life and the distribution of age-specific mortality and reproduction. Nature Ecology & Evolution. 3:1217–1224.

40. Hector J.A.L., Croxall J.P., Follett B.K. 1986. Reproductive endocrinology of the Wandering Albatross *Diomedea exulans* in relation to biennial breeding and deferred sexual maturity. Ibis. 128:9–22.

41. Heinsohn R., Cockburn A. 1997. Helping is costly to young birds in cooperatively breeding White-winged Choughs. Proceedings of the Royal Society B: Biological Sciences. 256:293–298.

42. Holekamp K.E., Strauss E.D. 2020. Reproduction within a hierarchical society from a female’s perspective. Integrative and Comparative Biology. 60:753–764.

43. Immelmann K. 1971. Ecological aspects of periodic reproduction. Avian Biology. Volume I.:341–398.

44. Koenig W.D., Pitelka F.A., Carmen W.J., Mumme R.L., Stanback M.T. 1992. The evolution of delayed dispersal in cooperative breeders. The Quarterly Review of Biology. 67:111– 150.

45. Kozłowski J. 1992. Optimal allocation of resources to growth and reproduction: Implications for age and size at maturity. Trends in Ecology & Evolution. 7:15–19.

46. Kralick A.E., O’Connell C.A., Bastian M.L., Hoke M.K., Zemel B.S., Schurr T.G., Tocheri M.W. 2023. Beyond dimorphism: Body size variation among adult orangutans is not dichotomous by sex. Integrative and Comparative Biology. 67:907–921.

47. Lack D. 1968. Ecological adaptations for breeding in birds. London: Methuen.

48. López-Calderón C., Hobson K.A., Marzal A., Balbontín J., Reviriego M., Magallanes S., García-Longoria L., de Lope F., Møller A.P. 2017. Wintering areas predict age-related breeding phenology in a migratory passerine bird. Journal of Avian Biology. 48:631–639.

49. Maddison W.P., FitzJohn R.G. 2015. The unsolved challenge to phylogenetic correlation tests for categorical characters. Systematic Biology. 64:127–136.

50. de Magalhães J.P., Costa J., Church G.M. 2007. An analysis of the relationship between metabolism, developmental schedules, and longevity using phylogenetic independent contrasts. The Journals of Gerontology A: Biological Sciences and Medical Sciences. 62:149–160.

51. Marchant S., Higgins P.J. 1990. The Handbook of Australian, New Zealand and Antarctic Birds. Melbourne: Oxford University Press.

52. McCormack J.E., Brown J.L. 2020. Mexican Jay (*Aphelocoma wollweberi*). In: Poole A.F., editor. Birds of the world. Ithaca: Cornell Lab of Ornithology.

53. McDonald D.B. 1989. Cooperation under sexual selection: Age-graded changes in a lekking bird. The American Naturalist. 134:709–730.

54. McDonald D.B. 1993. Demographic consequences of sexual selection in the Long-tailed Manakin. Behavioral Ecology. 4:297–309.

55. McTavish E.J., Gerbracht J.A., Holder M.T., Iliff M.J., Lepage D., Rasmussen P., Redelings B., Sanchez Reyes L.L., Miller E.T. 2024. A complete and dynamic tree of birds. bioRxiv. 10.1101/2024.05.20.595017.

56. Møller A.P. 2006. Sociality, age at first reproduction and senescence: Comparative analyses of birds. Journal of Evolutionary Biology. 19: 682–689.

57. Neate-Clegg M.H.C., Tingley M.W. 2023. Adult male birds advance spring migratory phenology faster than females and juveniles across North America. Global Change Biology. 29:341– 354.

58. Orians G.H. 1969. On the evolution of mating systems in birds and mammals. The American Naturalist. 103:589–603.

59. Pagel M. 1994. Detecting correlated evolution on phylogenies: a general method for the comparative analysis of discrete characters. Proceedings of the Royal Society B: Biological Sciences. 255:37–45.

60. Paradis E., Claude J., Strimmer K. 2004. APE: analyses of phylogenetics and evolution in R language. Bioinformatics. 20:289–290.

61. Parker P.G., Waite T.A., Decker M.D. 1995. Kinship and association in communally roosting Black Vultures. Animal Behaviour. 49:395–401.

62. Penndorf J., Ewart K.M., Klump B.C., Martin J.M., Aplin L.M. 2023. Social network analysis reveals context-dependent kin relationships in wild Sulphur-crested Cockatoos *Cacatua galerita*. Journal of Animal Ecology. 92:171–182.

63. Perrins C.M., Birkhead T.R. 1983. Social systems: Territoriality and coloniality. In: Avian Ecology. Glasgow: Blackie. p. 7–32.

64. Perry J.S. 1953. The reproduction of the African Elephant, *Loxodonta africana*. Philosophical Transactions of the Royal Society B: Biological Sciences. 643:93–149.

65. Pickering S.P.C. 1989. Attendance patterns and behaviour in relation to experience and pair-bond formation in the Wandering Albatross *Diomedea exulans* at South Georgia. Ibis. 131:183–195.

66. Pinheiro J.S., Bates D.M. 2000. Mixed-effects models in S and S-PLUS. New York: Springer-Verlag.

67. Prum R.O. 1994. Phylogenetic analysis of the evolution of alternative social behavior in the manakins (Aves: Pipridae). Evolution. 48:1657–1675.

68. Prum R.O. 2017. The evolution of beauty. New York: Anchor.

69. R Core Team. 2024. R: A language and environment for statistical computing.

70. Revell L.J. 2012. phytools: An R package for phylogenetic comparative biology (and other things). Methods in Ecology and Evolution. 3:217–223.

71. Riehl C. 2021. Evolutionary origins of cooperative and communal breeding: Lessons from the crotophagine cuckoos. Ethology. 127:827–836.

72. Riotte-Lambert L., Weimerskirch H. 2013. Do naive juvenile seabirds forage differently from adults? Proceedings of the Royal Society B: Biological Sciences. 280:20131434.

73. Robertson D.R. 1972. Social control of sex reversal in a coral-reef fish. Science. 177:1007–1009.

74. Rodd F.H., Reznick D.N., Sokolowski M.B. 1997. Phenotypic plasticity in the life history traits of guppies: Responses to social environment. Ecology. 78:419–433.

75. Roff D.A. 1984. The evolution of life history parameters in teleosts. Canadian Journal of Fisheries and Aquatic Sciences. 41:989–1000.

76. Rolland C., Danchin E., Fraipont M. de. 1998. The evolution of coloniality in birds in relation to food, habitat, predation, and life-history traits: A comparative analysis. The American Naturalist. 151:514–529.

77. Rowley I. 1978. Communal activities among White-winged Choughs *Corcorax melanorhamphus*. Ibis. 120:178–197.

78. Rubenstein D.R. 2011. Spatiotemporal environmental variation, risk aversion, and the evolution of cooperative breeding as a bet-hedging strategy. Proceedings of the National Academy of Sciences. 108:10816–10822.

79. Ryder T.B., Parker P.G., Blake J.G., Loiselle B.A. 2009. It takes two to tango: Reproductive skew and social correlates of male mating success in a lek-breeding bird. Proceedings of the Royal Society B: Biological Sciences. 276:2377–2384.

80. Salguero-Gómez R., Jones O.R., Jongejans E., Blomberg S.P., Hodgson D.J., Mbeau-Ache C., Zuidema P.A., De Kroon H., Buckley Y.M. 2016. Fast–slow continuum and reproductive strategies structure plant life-history variation worldwide. Proceedings of the National Academy of Sciences. 113:230–235.

81. Skutch A.F. 1961. Helpers among birds. The Condor. 63:198–226.

82. Slotow R., van Dyk G., Poole J., Page B., Klocke A. 2000. Older bull elephants control young males. Nature. 408:425–426.

83. Smith A.J., Robertson B.I. 1978. Social organization of Bell Miners. Emu. 78:169–178.

84. Snow D.W. 1971. Evolutionary aspects of fruit-eating by birds. Ibis. 113:194–202.

85. Stearns S.C. 1983. The influence of size and phylogeny on patterns of covariation among life-history traits in the mammals. Oikos. 41:173–187.

86. Stearns S.C. 1992. The evolution of life histories. New York: Oxford University Press.

87. Stearns S.C., Koella J.C. 1986. The evolution of phenotypic plasticity in life-history traits: predictions of reaction norms for age and size at maturity. Evolution. 40:893–913.

88. Taylor L.U. 2024. Young American Herring Gulls (*Larus argentatus* subsp. *smithsonianus*) have the opportunity for social development at the breeding colony. Waterbirds. 47:1–12.

89. Taylor L.U., Prum R.O. 2024. Developmental axioms in life history evolution. Biological Theory. 19:237–245.

90. Tobias J.A., Sheard C., Pigot A.L., Devenish A.J., Yang J., Sayol F., Neate-Clegg M.H., Alioravainen N., Weeks T.L., Barber R.A. 2022. AVONET: Morphological, ecological and geographical data for all birds. Ecology Letters. 25:581–597.

91. Trainer J.M., McDonald D.B., Learn W.A. 2002. The development of coordinated singing in cooperatively displaying long-tailed manakins. Behavioral Ecology. 13:65–69.

92. Uyeda J.C., Zenil-Ferguson R., Pennell M.W. 2018. Rethinking phylogenetic comparative methods. Systematic Biology. 67:1091–1109.

93. Vandenbergh J.G., Drickamer L.C., Colby D.R. 1972. Social and dietary factors in the sexual maturation of female mice. Reproduction. 28:397–405.

94. Weimerskirch H., Jouventin P. 1987. Population dynamics of the Wandering Albatross, *Diomedea exulans*, of the Crozet Islands: Causes and consequences of the population decline. Oikos. 49:315–322.

95. Welklin J.F., Lantz S.M., Khalil S., Moody N.M., Karubian J., Webster M.S. 2023. Photoperiod and rainfall are associated with seasonal shifts in social structure in a songbird. Behavioral Ecology. 34:136–149.

96. Western D., Ssemakula J. 1982. Life history patterns in birds and mammals and their evolutionary interpretation. Oecologia. 54:281–290.

97. Wickham H., Averick M., Bryan J., Chang W., McGowan L., François R., Grolemund G., Hayes A., Henry L., Hester J. 2019. Welcome to the tidyverse. Journal of Open Source Software. 4:1686.

98. Wikelski M., Hau M., Wingfield J.C. 2000. Seasonality of reproduction in a neotropical rain forest bird. Ecology. 81:2458–2472.

99. Wiley R.H. 1974. Evolution of social organization and life-history patterns among grouse. The Quarterly Review of Biology. 49:201–227.

100. Wiley R.H. 1978. The lek mating system of the Sage Grouse. Scientific American. 238:114–125.

101. Williams G.C. 1966. Adaptation and natural selection. Princeton: Princeton University Press.

102. Wingfield J.C., Farner D.S. 1980. Control of seasonal reproduction in temperate-zone birds. In: Reiter R.J., Follett B.K., editors. Seasonal reproduction in higher vertebrates. Basel: S. Karger. p. 62–101.

103. Wittenberger J.F. 1979. A model for delayed reproduction in iteroparous animals. The American Naturalist. 114:439–446.

104. Wunderle J. Joseph. 1991. Age-specific foraging proficiency in birds. Current Ornithology. 8:273–324.

105. Yu G., Smith D.K., Zhu H., Guan Y., Lam T.T.-Y. 2017. ggtree: An R package for visualization and annotation of phylogenetic trees with their covariates and other associated data. Methods in Ecology and Evolution. 8:28–36.

106. Zack S., Stutchbury B.J. 1992. Delayed breeding in avian social systems: The role of territory quality and “floater” tactics. Behaviour. 123:194–219.

