## Supplementary Material for "Social context and the evolution of delayed reproduction in birds"

**Table of Contents**

| | | |
|---|---|---|
| p. | 2 | Table S1 |
| p. | 3 | Table S2 |
| p. | 4 | Table S3 |
| pp. | 5–16 | Discrete model descriptions |

**Table S1**. Species in the dataset with observations of age at first breeding <1 yr. See main data file for references.

| **Order** | **Family** | **Taxon** | **Common** |
|---|---|---|---|
| Galliformes | Numididae | *Numida meleagris* | Helmeted Guineafowl |
| | Phasianidae | *Scleroptila afra* | Gray-winged Francolin |
| | | *Synoicus chinensis* | Blue-breasted Quail |
| | | *Coturnix coturnix* | Common Quail |
| | | *Coturnix pectoralis* | Stubble Quail |
| Columbiformes | Columbidae | *Columba livia* | Rock Pigeon |
| | | *Nesoenas mayeri* | Pink Pigeon |
| | | *Streptopelia decaocto* | Eurasian Collared-Dove |
| Caprimulgiformes | Trochilidae | *Calypte anna* | Anna's Hummingbird |
| Gruiformes | Rallidae | *Gallirallus australis* | Weka |
| Charadriiformes | Turnicidae | *Turnix sylvaticus* | Small Buttonquail |
| | | *Turnix velox* | Little Buttonquail |
| | | *Turnix melanogaster* | Black-breasted Buttonquail |
| Accipitriformes | Accipitridae | *Elanus scriptus* | Letter-winged Kite |
| Strigiformes | Tytonidae | *Tyto alba* | Barn Owl |
| Coliiformes | Coliidae | *Colius striatus* | Speckled Mousebird |
| Coraciiformes | Alcedinidae | *Corythornis cristatus* | Malachite Kingfisher |
| Psittaciformes | Psittaculidae | *Neophema bourkii* | Bourke's Parrot |
| | | *Neophema pulchella* | Turquoise Parrot |
| | | *Cyanoramphus auriceps* | Yellow-crowned Parakeet |
| | | *Melopsittacus undulatus* | Budgerigar |
| Passeriformes | Rhipiduridae | *Rhipidura fuliginosa* | New Zealand Fantail |
| | Petroicidae | *Petroica australis* | South Island Robin |
| | Chaetopidae | *Chaetops frenatus* | Cape Rockjumper |
| | Alaudidae | *Eremopterix leucotis* | Chestnut-backed Sparrow-Lark |
| | Panuridae | *Panurus biarmicus* | Bearded Reedling |
| | Cisticolidae | *Prinia gracilis* | Graceful Prinia |
| | | *Cisticola juncidis* | Zitting Cisticola |
| | Estrildidae | *Taeniopygia guttata* | Zebra Finch |
| | | *Mandingoa nitidula* | Green-backed Twinspot |
| | | *Estrilda astrild* | Common Waxbill |
| | | *Amadina fasciata* | Cut-throat |
| | | *Uraeginthus angolensis* | Southern Cordonbleu |
| | | *Lagonosticta senegala* | Red-billed Firefinch |
| | Passeridae | *Passer domesticus* | House Sparrow |
| | Fringillidae | *Loxia curvirostra* | Red Crossbill |
| | | *Loxia leucoptera* | White-winged Crossbill |

**Table S2.** Full phylogenetic regression estimates for age at first reproduction (AFR, $log_2$) in birds. Values show median across 100 trees, with 5–95% quantiles in parentheses.

| | | | **Dataset** | |
|---|---|---|---|---|
| **Model** | **Variable** | | **Female** | **Male** |
| Mass | Intercept | | 0.47 (-0.40, 0.79) | 0.27 (-0.79, 0.66) |
| | | *P* | *0.52 (0.31, 0.87)* | *0.56 (0.26, 0.83)* |
| | Mass ($log_{10}$) | | 0.08 (-0.02, 0.35) | 0.14 (0.03, 0.47) |
| | | *P* | *0.22 (<0.001, 0.91)* | *0.04 (<0.001, 0.69)* |
| Mass + Social | Intercept | | -0.20 (-0.52, 0.09) | -0.42 (-0.94, -0.10) |
| | | *P* | *0.80 (0.39, 0.98)* | *0.57 (0.15, 0.90)* |
| | Mass ($log_{10}$) | | 0.26 (0.18, 0.37) | 0.34 (0.24, 0.51) |
| | | *P* | *<0.001 (<0.001, 0.006)* | *<0.001 (<0.001, <0.001)* |
| | Cooperative | | 0.12 (0.06, 0.16) | 0.15 (0.12, 0.17) |
| | | *P* | *0.055 (0.004, 0.25)* | *0.03 (0.004, 0.08)* |
| | Colonial | | 0.17 (0.02, 0.27) | 0.20 (0.07, 0.31) |
| | | *P* | *<0.001 (<0.001, 0.46)* | *<0.001 (<0.001, 0.02)* |
| | Lekking | | 0.15 (0.09, 0.22) | 0.62 (0.52, 0.73) |
| | | *P* | *0.53 (0.40, 0.69)* | *0.02 (0.002, 0.04)* |
| Mass * Social | Intercept | | -0.13 (-0.50, 0.15) | -0.44 (-0.97, -0.11) |
| | | *P* | *0.84 (0.42, 0.99)* | *0.57 (0.13, 0.90)* |
| | Mass ($log_{10}$) | | 0.25 (0.15, 0.36) | 0.34 (0.23, 0.50) |
| | | *P* | *<0.001 (<0.001, 0.03)* | *<0.001 (<0.001, 0.001)* |
| | Cooperative | | -0.01 (-0.09, 0.14) | -0.10 (-0.15, -0.02) |
| | | *P* | *0.80 (0.45, 0.96)* | *0.62 (0.46, 0.87)* |
| | Colonial | | -0.15 (-0.48, 0.03) | 0.05 (-0.30, 0.28) |
| | | *P* | *0.40 (0.03, 0.94)* | *0.56 (0.08, 0.96)* |
| | Lekking | | 0.03 (-0.09, 0.19) | 1.13 (0.86, 1.41) |
| | | *P* | *0.93 (0.77, 0.99)* | *0.17 (0.04, 0.34)* |
| | Mass * Cooperative | | 0.06 (-0.03, 0.11) | 0.11 (0.06, 0.15) |
| | | *P* | *0.46 (0.24, 0.94)* | *0.19 (0.10, 0.47)* |
| | Mass * Colonial | | 0.09 (0.002, 0.24) | 0.04 (-0.06, 0.19) |
| | | *P* | *0.13 (0.001, 0.90)* | *0.42 (0.01, 0.89)* |
| | Mass * Lekking | | 0.05 (-0.03, 0.11) | -0.18 (-0.30, -0.07) |
| | | *P* | *0.85 (0.69, 0.98)* | *0.55 (0.21, 0.83)* |

**Table S3**. Continuous evolutionary model estimates for age at first reproduction (AFR, $log_2$) in birds. Values show median across 100 trees, with 5–95% quantiles in parentheses.

| | | **Dataset** | |
|---|---|---|---|
| **Model** | **Variable** | **Female** | **Male** |
| BM | $\sigma^2$ | 0.043 (0.022, 0.049) | 0.048 (0.028, 0.054) |
| OU1 | $\alpha$ | 0.056 (0.025, 0.066) | 0.060 (0.030, 0.069) |
| | $\sigma^2$ | 0.068 (0.029, 0.081) | 0.079 (0.039, 0.093) |
| | $\theta$ | 0.645 (0.615, 0.679) | 0.697 (0.670, 0.729) |
| OU2-Cooperative | $\alpha$ | 0.056 (0.025, 0.066) | 0.060 (0.030, 0.069) |
| | $\sigma^2$ | 0.068 (0.030, 0.082) | 0.079 (0.039, 0.094) |
| | $\theta$, Cooperative | 0.598 (0.517, 0.787) | 0.582 (0.488, 0.674) |
| | $\theta$, Everything else | 0.646 (0.623, 0.687) | 0.708 (0.684, 0.743) |
| OU2-Colonial | $\alpha$ | 0.078 (0.029, 0.095) | 0.079 (0.035, 0.095) |
| | $\sigma^2$ | 0.079 (0.031, 0.096) | 0.091 (0.041, 0.108) |
| | $\theta$, Colonial | 1.254 (1.185, 1.371) | 1.283 (1.221, 1.371) |
| | $\theta$, Everything else | 0.455 (0.417, 0.536) | 0.519 (0.482, 0.575) |
| OU2-Lekking | $\alpha$ | 0.056 (0.025, 0.065) | 0.061 (0.032, 0.071) |
| | $\sigma^2$ | 0.068 (0.029, 0.081) | 0.078 (0.038, 0.092) |
| | $\theta$, Lekking | 1.102 (0.902, 1.856) | 2.272 (1.966, 3.310) |
| | $\theta$, Everything else | 0.632 (0.607, 0.667) | 0.650 (0.621, 0.683) |
| OU2-Social | $\alpha$ | 0.066 (0.028, 0.078) | 0.074 (0.035, 0.087) |
| | $\sigma^2$ | 0.073 (0.030, 0.087) | 0.087 (0.041, 0.101) |
| | $\theta$, Social | 1.031 (0.965, 1.194) | 1.128 (1.061, 1.283) |
| | $\theta$, Other | 0.433 (0.396, 0.488) | 0.449 (0.413, 0.487) |
| OU4-Social | $\alpha$ | 0.078 (0.030, 0.095) | 0.085 (0.039, 0.102) |
| | $\sigma^2$ | 0.079 (0.031, 0.096) | 0.092 (0.041, 0.109) |
| | $\theta$, Cooperative | 0.563 (0.486, 0.772) | 0.543 (0.493, 0.698) |
| | $\theta$, Colonial | 1.257 (1.186, 1.380) | 1.284 (1.224, 1.358) |
| | $\theta$, Lekking | 0.940 (0.780, 1.663) | 1.953 (1.739, 2.992) |
| | $\theta$, Other | 0.414 (0.379, 0.482) | 0.431 (0.395, 0.481) |

**Discrete model descriptions**

We compared 11 discrete models to test the evolutionary relationship between delayed reproduction and social context in birds. Species were binned into two age at first reproduction (AFR) states ("Fast" = AFR ≤ 2 *vs.* "Slow" = AFR ≥ 3) and two social context states ("Social" = cooperative, colonial, or lekking species *vs*. "Other"). More complicated models included a hidden character, the state of which (H1 or H2) was identifiable insofar as it affects transition rate parameters among observed character states.

For each model, we first present the model structure for transition rate parameters. Individual parameters are given as italicized capital letters. There is no relationship between parameters from different models. Excluded transitions (i.e., constrained at rate 0) are left blank. All models were asymmetrical (e.g., different parameters for Fast/Social → Fast/Other *vs*. Fast/Other → Fast/Social) and excluded non-stepwise transitions (e.g., Fast/Social → Slow/Other). Following each model structure, we list the empirical transition rate parameters (median values across 100 trees, with 5–95% quantiles in parentheses).

Note that simpler models (models #1–4) could also be shown with a hidden state difference (H1 or H2). We collapse these models because no parameters differed with respect to hidden state, and the hidden state parameters were therefore unidentifiable. However, such an expanded view would highlight how H1 dependent and H2 independent models are complementary. For example, model #2 relates AFR transitions to social state, whereas model #5 relates AFR transitions to hidden state, the difference being we provide data on social but not hidden states.

**Model 1: H1-Independent**

Transitions between AFR states (Fast ↔ Slow) are independent of social state.

Transitions between social states (Other ↔ Social) are independent of AFR state.

| | **Fast Other** | **Fast Social** | **Slow Other** | **Slow Social** |
|---|---|---|---|---|
| **Fast Other** | - | *A* | *C* | |
| **Fast Social** | *B* | - | | *C* |
| **Slow Other** | *D* | | - | *A* |
| **Slow Social** | | *D* | *B* | - |

| **Parameter** | **Description** | **Female estimate** | **Male estimate** |
|---|---|---|---|
| A | Other → Social | 0.0177 (0.0170, 0.0183) | 0.0178 (0.0171, 0.0184) |
| B | Social → Other | 0.0135 (0.0110, 0.0180) | 0.0140 (0.0111, 0.0181) |
| C | Fast → Slow | 0.0029 (0.0026, 0.0031) | 0.0042 (0.0039, 0.0045) |
| D | Slow → Fast | 0.0253 (0.0239, 0.0269) | 0.0251 (0.0238, 0.0266) |

**<u>Model 2: H1-Dependent-AFR</u>**

Transitions between AFR states (Fast ↔ Slow) differ by social state.

Transitions between social states (Other ↔ Social) are independent of AFR state.

| | **Fast Other** | **Fast Social** | **Slow Other** | **Slow Social** |
|---|---|---|---|---|
| **Fast Other** | - | *A* | *C* | |
| **Fast Social** | *B* | - | | *D* |
| **Slow Other** | *E* | | - | *A* |
| **Slow Social** | | *F* | *B* | - |

| **Parameter** | **Description** | **Female estimate** | **Male estimate** |
|---|---|---|---|
| A | Other → Social | 0.0181 (0.0174, 0.0188) | 0.0182 (0.0176, 0.0189) |
| B | Social → Other | 0.0131 (0.0109, 0.0169) | 0.0128 (0.0107, 0.0162) |
| C | Fast → Slow, Other | 0.0011 (0.0009, 0.0014) | 0.0018 (0.0014, 0.0021) |
| D | Fast → Slow, Social | 0.0133 (0.0112, 0.0142) | 0.0179 (0.0164, 0.0191) |
| E | Slow → Fast, Other | 0.0351 (0.0318, 0.0401) | 0.0347 (0.0320, 0.0387) |
| F | Slow → Fast, Social | 0.0162 (0.0132, 0.0204) | 0.0178 (0.0151, 0.0202) |

**Model 3: H1-Dependent-Social**

Transitions between AFR states (Fast ↔ Slow) are independent of social state.

Transitions between social states (Other ↔ Social) differ by AFR state.

| | **Fast Other** | **Fast Social** | **Slow Other** | **Slow Social** |
|---|---|---|---|---|
| **Fast Other** | - | *A* | *E* | |
| **Fast Social** | *C* | - | | *E* |
| **Slow Other** | *F* | | - | *B* |
| **Slow Social** | | *F* | *D* | - |

| **Parameter** | **Description** | **Female estimate** | **Male estimate** |
|---|---|---|---|
| A | Other → Social, Fast | 0.0177 (0.0171, 0.0184) | 0.0173 (0.0166, 0.0181) |
| B | Other → Social, Slow | 0.0193 (0.0170, 0.0220) | 0.0228 (0.0207, 0.0269) |
| C | Social → Other, Fast | 0.0385 (0.0359, 0.0407) | 0.0408 (0.0386, 0.0439) |
| D | Social → Other, Slow | 0.0011 (0.0000, 0.0016) | 0.0009 (0.0000, 0.0014) |
| E | Fast → Slow | 0.0030 (0.0028, 0.0033) | 0.0045 (0.0042, 0.0053) |
| F | Slow → Fast | 0.0259 (0.0240, 0.0275) | 0.0253 (0.0219, 0.0270) |

**Model 4: H1-Dependent-Both**

Transitions between AFR states (Fast ↔ Slow) differ by social state.

Transitions between social states (Other ↔ Social) differ by AFR state.

| | **Fast Other** | **Fast Social** | **Slow Other** | **Slow Social** |
|---|---|---|---|---|
| **Fast Other** | - | *A* | *E* | |
| **Fast Social** | *C* | - | | *F* |
| **Slow Other** | *G* | | - | *B* |
| **Slow Social** | | *H* | *D* | - |

| Parameter | Description | Female estimate | Male estimate |
|---|---|---|---|
| A | Other → Social, Fast | 0.0181 (0.0172, 0.0191) | 0.0177 (0.0165, 0.0189) |
| B | Other → Social, Slow | 0.0166 (0.0141, 0.0221) | 0.0212 (0.0156, 0.0346) |
| C | Social → Other, Fast | 0.0373 (0.0342, 0.0400) | 0.0399 (0.0374, 0.0431) |
| D | Social → Other, Slow | 0.0014 (0.0000, 0.0018) | 0.0008 (0.0000, 0.0016) |
| E | Fast → Slow, Other | 0.0023 (0.0019, 0.0026) | 0.0036 (0.0029, 0.0051) |
| F | Fast → Slow, Social | 0.0048 (0.0040, 0.0058) | 0.0065 (0.0052, 0.0083) |
| G | Slow → Fast, Other | 0.0273 (0.0204, 0.0310) | 0.0234 (0.0083, 0.0289) |
| H | Slow → Fast, Social | 0.0249 (0.0218, 0.0304) | 0.0262 (0.0229, 0.0305) |

**Model 5: H2-Independent-AFR**

Transitions between AFR states (Fast ↔ Slow) differ by hidden state.

Transitions between social states (Other ↔ Social) are independent.

Transitions between hidden states (H1 ↔ H2) are independent.

| | **H1** Fast Other | **H1** Fast Social | **H1** Slow Other | **H1** Slow Social | **H2** Fast Other | **H2** Fast Social | **H2** Slow Other | **H2** Slow Social |
|---|---|---|---|---|---|---|---|---|
| **H1 Fast Other** | - | *A* | *C* | | *G* | | | |
| **H1 Fast Social** | *B* | - | | *C* | | *G* | | |
| **H1 Slow Other** | *E* | | - | *A* | | | *G* | |
| **H1 Slow Social** | | *E* | *B* | - | | | | *G* |
| **H2 Fast Other** | *G* | | | | - | *A* | *D* | |
| **H2 Fast Social** | | *G* | | | *B* | - | | *D* |
| **H2 Slow Other** | | | *G* | | *F* | | - | *A* |
| **H2 Slow Social** | | | | *G* | | *F* | *B* | - |

| Parameter | Description | Female estimate | Male estimate |
|---|---|---|---|
| A | Other → Social | 0.0177 (0.017,0 0.0183) | 0.0178 (0.0171, 0.0184) |
| B | Social → Other | 0.0135 (0.0110, 0.0180) | 0.0140 (0.0111, 0.0181) |
| C | Fast → Slow, H1 | 0.0000 (0.0000, 0.0526) | 0.0016 (0.0000, 0.136) |
| D | Fast → Slow, H2 | 0.0449 (0.0003, 0.0609) | 0.0614 (0.0000, 0.0995) |
| E | Slow → Fast, H1 | 0.1119 (0.0155, 0.2789) | 0.1439 (0.0064, 19.2445) |
| F | Slow → Fast, H2 | 0.0184 (0.0144, 0.1100) | 0.0190 (0.0110, 0.1378) |
| G | H1 ↔ H2 | 0.0058 (0.0048, 0.0063) | 0.0068 (0.0051, 0.0082) |

**Model 6: H2-Independent-Social**

Transitions between AFR states (Fast ↔ Slow) are independent.

Transitions between social states (Other ↔ Social) differ by hidden state.

Transitions between hidden states (H1 ↔ H2) are independent.

| | **H1 Fast Other** | **H1 Fast Social** | **H1 Slow Other** | **H1 Slow Social** | **H2 Fast Other** | **H2 Fast Social** | **H2 Slow Other** | **H2 Slow Social** |
|---|---|---|---|---|---|---|---|---|
| **H1 Fast Other** | - | *A* | *E* | | *G* | | | |
| **H1 Fast Social** | *B* | - | | *E* | | *G* | | |
| **H1 Slow Other** | *F* | | - | *A* | | | *G* | |
| **H1 Slow Social** | | *F* | *B* | - | | | | *G* |
| **H2 Fast Other** | *G* | | | | - | *C* | *E* | |
| **H2 Fast Social** | | *G* | | | *D* | - | | *E* |
| **H2 Slow Other** | | | *G* | | *F* | | - | *C* |
| **H2 Slow Social** | | | | *G* | | *F* | *D* | - |

| Parameter | Description | Female estimate | Male estimate |
|---|---|---|---|
| A | Other → Social, H1 | 7.2623 (0.0047, 37.5839) | 7.2890 (0.0047, 79.1381) |
| B | Social → Other, H1 | 10.1082 (0.0000, 43.7577) | 9.2124 (0.0000, 100[MAX]) |
| C | Other → Social, H2 | 0.0073 (0.0043, 80.7865) | 0.0079 (0.0006, 15.3267) |
| D | Social → Other, H2 | 0.0000 (0.0000, 100[MAX]) | 0.0000 (0.0000, 43.6459) |
| E | Fast → Slow | 0.0029 (0.0026, 0.0031) | 0.0042 (0.0039, 0.0045) |
| F | Slow → Fast | 0.0253 (0.0239, 0.0269) | 0.0251 (0.0238, 0.0266) |
| G | H1 ↔ H2 | 0.0100 (0.0085, 0.0115) | 0.0094 (0.0082, 0.0128) |

**Model 7: H2-Dependent-AFR**

Transitions between AFR states (Fast ↔ Slow) differ by social and hidden states.

Transitions between social states (Other ↔ Social) are independent.

Transitions between hidden states (H1 ↔ H2) are independent.

| | **H1** Fast Other | **H1** Fast Social | **H1** Slow Other | **H1** Slow Social | **H2** Fast Other | **H2** Fast Social | **H2** Slow Other | **H2** Slow Social |
|---|---|---|---|---|---|---|---|---|
| **H1 Fast Other** | - | *A* | *C* | | *K* | | | |
| **H1 Fast Social** | *B* | - | | *D* | | *K* | | |
| **H1 Slow Other** | *E* | | - | *A* | | | *K* | |
| **H1 Slow Social** | | *F* | *B* | - | | | | *K* |
| **H2 Fast Other** | *K* | | | | - | *A* | *G* | |
| **H2 Fast Social** | | *K* | | | *B* | - | | *H* |
| **H2 Slow Other** | | | *K* | | *I* | | - | *A* |
| **H2 Slow Social** | | | | *K* | | *J* | *B* | - |

| Parameter | Description | Female estimate | Male estimate |
|---|---|---|---|
| A | Other → Social | 0.0179 (0.0173, 0.0185) | 0.0180 (0.0174, 0.0186) |
| B | Social → Other | 0.0123 (0.0103, 0.0148) | 0.0122 (0.0102, 0.0143) |
| C | Fast → Slow, Other/H1 | 0.0434 (0.0050, 0.0675) | 0.0346 (0.0121, 0.0504) |
| D | Fast → Slow, Social/H1 | 0.0688 (0.0604, 0.0780) | 0.0909 (0.0797, 0.1070) |
| E | Slow → Fast, Other/H1 | 0.4093 (0.0236, 0.6619) | 0.4906 (0.1404, 0.7276) |
| F | Slow → Fast, Social/H1 | 0.0941 (0.0730, 0.1119) | 0.1092 (0.0913, 0.1294) |
| G | Fast → Slow, Other/H2 | 0.0000 (0.0000, 0.0000) | 0.0000 (0.0000, 0.0000) |
| H | Fast → Slow, Social/H2 | 0.0000 (0.0000, 0.0000) | 0.0000 (0.0000, 0.0013) |
| I | Slow → Fast, Other/H2 | 0.0284 (0.0235, 0.0429) | 0.0255 (0.0224, 0.0286) |
| J | Slow → Fast, Social/H2 | 0.0066 (0.0036, 0.0082) | 0.0065 (0.0040, 0.0080) |
| K | H1 ↔ H2 | 0.0061 (0.0052, 0.0084) | 0.0070 (0.0063, 0.0085) |

**Model 8: H2-Dependent-Social**

Transitions between AFR states (Fast ↔ Slow) are independent.

Transitions between social states (Other ↔ Social) differ by AFR and hidden states.

Transitions between hidden states (H1 ↔ H2) are independent.

| | **H1 Fast Other** | **H1 Fast Social** | **H1 Slow Other** | **H1 Slow Social** | **H2 Fast Other** | **H2 Fast Social** | **H2 Slow Other** | **H2 Slow Social** |
|---|---|---|---|---|---|---|---|---|
| **H1 Fast Other** | - | *A* | *I* | | *K* | | | |
| **H1 Fast Social** | *C* | - | | *I* | | *K* | | |
| **H1 Slow Other** | *J* | | - | *B* | | | *K* | |
| **H1 Slow Social** | | *J* | *D* | - | | | | *K* |
| **H2 Fast Other** | *K* | | | | - | *E* | *I* | |
| **H2 Fast Social** | | *K* | | | *G* | - | | *I* |
| **H2 Slow Other** | | | *K* | | *J* | | - | *F* |
| **H2 Slow Social** | | | | *K* | | *J* | *H* | - |

| Parameter | Description | Female estimate | Male estimate |
|---|---|---|---|
| A | Other → Social, Fast/H1 | 10.7281 (0.0588, 75.7751) | 5.9650 (0.0300, 33.7045) |
| B | Other → Social, Slow/H1 | 0.2313 (0.0172, 0.5599) | 0.2972 (0.0717, 100[MAX]) |
| C | Social → Other, Fast/H1 | 18.0260 (0.0990, 100[MAX]) | 21.3576 (0.0294, 100[MAX]) |
| D | Social → Other, Slow/H1 | 0.2515 (0.0000, 1.7110) | 0.6966 (0.0000, 100[MAX]) |
| E | Other → Social, Fast/H2 | 0.0056 (0.0012, 0.5718) | 0.0013 (0.0005, 2.0197) |
| F | Other → Social, Slow/H2 | 0.0183 (0.0142, 0.3420) | 0.0690 (0.0174, 100[MAX]) |
| G | Social → Other, Fast/H2 | 0.0191 (0.0151, 0.7426) | 0.0225 (0.0086, 17.7547) |
| H | Social → Other, Slow/H2 | 0.0000 (0.0000, 0.0013) | 0.0000 (0.0000, 0.8124) |
| I | Fast → Slow | 0.0029 (0.0027, 0.0032) | 0.0044 (0.0041, 0.0046) |
| J | Slow → Fast | 0.0256 (0.0241, 0.0273) | 0.0251 (0.0235, 0.0267) |
| K | H1 ↔ H2 | 0.0071 (0.0059, 0.0086) | 0.0074 (0.0060, 0.0089) |

**Model 9: H2-Dependent-AFR-Hidden-Social**

Transitions between AFR states (Fast ↔ Slow) differ by social state.

Transitions between social states (Other ↔ Social) differ by hidden state.

Transitions between hidden states (H1 ↔ H2) are independent.

| | **H1** Fast Other | **H1** Fast Social | **H1** Slow Other | **H1** Slow Social | **H2** Fast Other | **H2** Fast Social | **H2** Slow Other | **H2** Slow Social |
|---|---|---|---|---|---|---|---|---|
| **H1 Fast Other** | - | *A* | *E* | | *I* | | | |
| **H1 Fast Social** | *C* | - | | *F* | | *I* | | |
| **H1 Slow Other** | *G* | | - | *A* | | | *I* | |
| **H1 Slow Social** | | *H* | *C* | - | | | | *I* |
| **H2 Fast Other** | *I* | | | | - | *B* | *E* | |
| **H2 Fast Social** | | *I* | | | *D* | - | | *F* |
| **H2 Slow Other** | | | *I* | | *G* | | - | *B* |
| **H2 Slow Social** | | | | *I* | | *H* | *D* | - |

| Parameter | Description | Female estimate | Male estimate |
|---|---|---|---|
| A | Other → Social, H1 | 7.1751 (0.0052, 31.2181) | 1.5182 (0.0073, 17.5938) |
| B | Other → Social, H2 | 0.0080 (0.0014, 76.6122) | 0.0281 (0.0073, 73.4938) |
| C | Social → Other, H1 | 10.5070 (0.0000, 80.0110) | 6.4157 (0.0000, 39.542) |
| D | Social → Other, H2 | 0.0000 (0.0000, 100[MAX]) | 0.0000 (0.0000, 100[MAX]) |
| E | Fast → Slow, Other | 0.0005 (0.0000, 0.0008) | 0.0012 (0.0000, 0.0016) |
| F | Fast → Slow, Social | 0.0163 (0.0142, 0.0190) | 0.0220 (0.0193, 0.0276) |
| G | Slow → Fast, Other | 0.0382 (0.0337, 0.0549) | 0.0366 (0.0251, 0.0422) |
| H | Slow → Fast, Social | 0.0138 (0.0111, 0.0172) | 0.0160 (0.0134, 0.0191) |
| I | H1 ↔ H2 | 0.0099 (0.0081, 0.0141) | 0.0083 (0.0061, 0.0100) |

**Model 10: H2-Dependent-Social-Hidden-AFR**

Transitions between AFR states (Fast ↔ Slow) differ by hidden state.

Transitions between social states (Other ↔ Social) differ by AFR state.

Transitions between hidden states (H1 ↔ H2) are independent.

| | H1 Fast Other | H1 Fast Social | H1 Slow Other | H1 Slow Social | H2 Fast Other | H2 Fast Social | H2 Slow Other | H2 Slow Social |
|---|---|---|---|---|---|---|---|---|
| **H1 Fast Other** | - | *A* | *E* | | *I* | | | |
| **H1 Fast Social** | *C* | - | | *E* | | *I* | | |
| **H1 Slow Other** | *G* | | - | *B* | | | *I* | |
| **H1 Slow Social** | | *G* | *D* | - | | | | *I* |
| **H2 Fast Other** | *I* | | | | - | *A* | *F* | |
| **H2 Fast Social** | | *I* | | | *C* | - | | *F* |
| **H2 Slow Other** | | | *I* | | *H* | | - | *B* |
| **H2 Slow Social** | | | | *I* | | *H* | *D* | - |

| Parameter | Description | Female estimate | Male estimate |
|---|---|---|---|
| A | Other → Social, Fast | 0.0168 (0.0161, 0.0176) | 0.0164 (0.0157, 0.0172) |
| B | Other → Social, Slow | 0.0293 (0.0253, 0.0387) | 0.0370 (0.0309, 0.0433) |
| C | Social → Other, Fast | 0.0354 (0.0333, 0.0374) | 0.0372 (0.0350, 0.0396) |
| D | Social → Other, Slow | 0.0000 (0.0000, 0.0000) | 0.0000 (0.0000, 0.0000) |
| E | Fast → Slow, H1 | 0.0000 (0.0000, 0.0439) | 0.0035 (0.0000, 0.1218) |
| F | Fast → Slow, H2 | 0.0574 (0.0331, 0.0969) | 0.0744 (0.0000, 0.1183) |
| G | Slow → Fast, H1 | 0.1033 (0.0041, 0.1644) | 0.0902 (0.0050, 1.1224) |
| H | Slow → Fast, H2 | 0.0234 (0.0151, 0.0748) | 0.0231 (0.0023, 0.0889) |
| I | H1 ↔ H2 | 0.0057 (0.0052, 0.0063) | 0.0072 (0.0060, 0.0078) |

**Model 11: H2-Dependent-Both**

Transitions between AFR states (Fast ↔ Slow) differ by social and hidden states.

Transitions between social states (Other ↔ Social) differ by AFR and hidden states.

Transitions between hidden states (H1 ↔ H2) are independent.

| | **H1** **Fast Other** | **H1** **Fast Social** | **H1** **Slow Other** | **H1** **Slow Social** | **H2** **Fast Other** | **H2** **Fast Social** | **H2** **Slow Other** | **H2** **Slow Social** |
|---|---|---|---|---|---|---|---|---|
| **H1 Fast Other** | - | *A* | *I* | | *Q* | | | |
| **H1 Fast Social** | *C* | - | | *J* | | *Q* | | |
| **H1 Slow Other** | *K* | | - | *B* | | | *Q* | |
| **H1 Slow Social** | | *L* | *D* | - | | | | *Q* |
| **H2 Fast Other** | *Q* | | | | - | *E* | *M* | |
| **H2 Fast Social** | | *Q* | | | *G* | - | | *N* |
| **H2 Slow Other** | | | *Q* | | *O* | | - | *F* |
| **H2 Slow Social** | | | | *Q* | | *P* | *H* | - |

| **Parameter** | **Description** | **Female estimate** | **Male estimate** |
|---|---|---|---|
| A | Other → Social, Fast/H1 | 0.0869 (0.0000, 0.1120) | 0.0388 (0.0000, 0.0430) |
| B | Other → Social, Slow/H1 | 0.0000 (0.0000, 1.2755) | 0.4052 (0.0750, 0.8939) |
| C | Social → Other, Fast/H1 | 0.0000 (0.0000, 0.0539) | 0.0493 (0.0000, 0.0555) |
| D | Social → Other, Slow/H1 | 0.0014 (0.0000, 3.6056) | 1.3081 (0.0000, 2.8836) |
| E | Other → Social, Fast/H2 | 0.0159 (0.0000, 19.5135) | 0.0000 (0.0000, 0.0429) |
| F | Other → Social, Slow/H2 | 0.3265 (0.0000, 0.9286) | 0.0927 (0.0726, 0.9898) |
| G | Social → Other, Fast/H2 | 0.0344 (0.0000, 42.8427) | 0.0000 (0.0000, 0.0555) |
| H | Social → Other, Slow/H2 | 1.2000 (0.0000, 3.6109) | 0.0002 (0.0000, 3.3312) |
| I | Fast → Slow, Other/H1 | 0.0243 (0.0002, 1.5694) | 0.0000 (0.0000, 3.8713) |
| J | Fast → Slow, Social/H1 | 0.0504 (0.0005, 0.0624) | 0.0019 (0.0012, 0.0850) |
| K | Slow → Fast, Other/H1 | 0.3354 (0.0204, 100[MAX]) | 0.0305 (0.0199, 100[MAX]) |
| L | Slow → Fast, Social/H1 | 0.0188 (0.0000, 0.0924) | 0.0247 (0.0000, 0.0370) |
| M | Fast → Slow, Other/H2 | 0.0008 (0.0002, 0.6748) | 0.4137 (0.0000, 4.0922) |
| N | Fast → Slow, Social/H2 | 0.0000 (0.0000, 0.0627) | 0.0707 (0.0012, 0.0871) |
| O | Slow → Fast, Other/H2 | 0.0361 (0.0131, 45.0391) | 11.8766 (0.0217, 100[MAX]) |
| P | Slow → Fast, Social/H2 | 0.0057 (0.0000, 0.0424) | 0.0268 (0.0000, 0.0368) |
| Q | H1 ↔ H2 | 0.0039 (0.0032, 0.0160) | 0.0073 (0.0067, 0.0078) |
